# Host identity and functional traits determine the community composition of the arbuscular mycorrhizal fungi in facultative epiphytic plant species

**DOI:** 10.1101/307991

**Authors:** MM Alguacil, G Díaz, P Torres, G Rodríguez-Caballero, A Roldán

**Affiliations:** Soil and Water Conservation Department, Centro de Edafología y Biología Aplicada del Segura-Consejo Superior de Investigaciones Científicas (CEBAS-CSIC), Campus Universitario de Espinardo, Murcia, Spain.; Departamento de Biología Aplicada. Área de Botánica. Universidad Miguel Hernández. Avda. De la Universidad s/n. 03202-Elche, Alicante, Spain.

**Keywords:** Facultative epiphytes, arbuscular mycorrhizal fungi, diversity, SSU rDNA, semiarid ecosystems

## Abstract

The epiphytic vascular flora is scarce and facultative in semiarid Mediterranean ecosystems, thus covering diverse taxonomic groups. However, differently to terrestrial conditions, little is known about the factors driving mycorrhizal communities in epiphytic environments. Here, we investigated the arbuscular mycorrhizal fungi (AMF) harboured by 31 plant species occurring in the trunks of *Phoenix dactylifera*. We wanted to ascertain if host identity and plant functional traits shape mycorrhizal communities. Specifically, we tested the plant life-cycle (perennial versus annual), the plant life-form (herbaceous versus woody), the plant origin (exotic versus native) and the plant species.

The roots were examined by molecular and phylogenetic analysis of AMF community. The plant affiliation to species strongly influenced the AMF assemblages. Plant life-form and plant life-cycle also shaped AMF interactions. The AMF community differed between annual and perennial species and higher AMF richness was detected in perennial plants. The indicator species analysis revealed three Operational Taxonomic Units belonging to the *Glomeraceae*, associated with annual species. However, the epiphytic plants associated with AMF irrespective of whether they were native or not, probably because here no functional differences derive from plant origin.

**IMPORTANCE:** Arbuscular mycorrhizal (AM) symbiosis has a decisive role in plant nutrient and water uptake by plants, with particular importance in stressful environments. Under semiarid conditions, the facultative epiphytic flora should cope with harsh conditions. While numerous studies have been conducted on factors driving terrestrial AM assemblages, the epiphytic environment remains unexplored. We offer new insights into composition of AM communities as shaped by epiphytic plant host identity and functional traits.

## INTRODUCTION

Epiphytic habitats are considered as extreme plant environments due to the large fluxes of temperature and low water and nutrient availability they are subjected to. In these conditions, symbiotic associations such as mycorrhizal symbiosis could be crucial because of their widely demonstrated role in plant nutrient and water uptake in terrestrial habits, with particular importance in stressful environments (38). Furthermore, the availability of compatible and suitable mycorrhizal fungi might be a key factor constraining the development and distribution of epiphytic plants and *vice versa*.

In semiarid Mediterranean ecosystems the epiphytic vascular flora is scarce and facultative or accidental, mainly occurring on the trunks of certain palm species. Here, the microhabitat conditions that originate in the cut leaves formed by the pruning of dead or old leaves allow water and debris accumulation, which enables occasional plant establishment (44,52).

Since most vascular epiphytic plants are ferns and their relatives or monocots (11), the mycorrhizal condition, specificity, and dependence have therefore mainly been studied in these taxonomic groups. Most studies concerning epiphytism-mycorrhizas relationships have been conducted in humid tropical habitats and involve orchids, which form mycorrhizas with Basidiomycetes (23,30,31), or other species that form arbuscular mycorrhizas, like those of the Araceae, Clusiaceae, Bromeliaceae, or Begoniaceae (20,33,36). However, very little is known about the facultative epiphytic plant species, belonging to diverse phylogenetic groups, which grow under semiarid Mediterranean conditions. Only the widespread *Sonchus tenerrimus* has been reported to establish a symbiotic relationship with arbuscular mycorrhizal fungi (AMF), belonging to the *Glomeromycota* (44).

But more interesting than checking the mycorrhizal status of epiphytic plants is to understand the factors driving symbiotic communities in epiphytic habitats and consequently their role in the tree canopy ecosystems, which still has been investigated less than in their terrestrial counterparts.

There is evidence that AMF communities in terrestrial habitats depend on host-plant preference (1,22,42), but the specificity in biotic interactions has been suggested to be mediated by plant functional traits rather than phylogeny (19,37). Plant life-cycle, for instance, may explain differences in AMF communities (2,41). It is likely that differences in AMF communities are due not only to plant life-cycle but also to a combination of factors such as life-form, physiology, phylogeny, or host origin (native vs exotic). It is expected that native plant species have co-evolved together with local AMF and thus their mycorrhizal communities would be different and more diverse to those of exotic plants in a particular environment. Some studies have demonstrated that the AMF associated with exotic plant invaders shift in both abundance and community composition, compared with native species (4,5,13,26). Moreover, functional traits are often conserved during evolution, resulting in closely related species that tend to interact with similar species in such a way that significant correlations have been observed between the phylogenetic composition of plants and AMF assemblages (24). But, as previous reports show, the AMF communities on epiphytic plants are different from those of the surrounding soil habitats (23,44). These results suggest that facultative epiphytic plants from semiarid sites might support distinctive AMF communities shaped by particular plant traits.

In this study we investigated facultative epiphytic plant species occurring in the trunks of *Phoenix dactylifera* trees cultivated in orchards under Mediterranean semiarid conditions and we assessed the arbuscular mycorrhizal communities they harbored. We wanted to ascertain 1) if there are AMF in plants growing epiphytically and 2) if host identity and plant functional traits shape these mycorrhizal communities. Specifically, we tested the plant life-cycle (perennial versus annual), the plant life-form (herbaceous versus woody), the plant origin (exotic versus native) and the plant species as driving factors.

## RESULTS

### PCR and sequence analysis

Arbuscular mycorrhizal fungal DNA was successfully amplified from 93 root samples (corresponding to 31 facultative epiphytic species) (Table 1) by nested PCR with the primer combination AML1/AML2, and generated PCR products of the expected band of approximately 795 bps, which were used for cloning and creating the clone libraries. Ninety-three clone libraries, were created. We screened 2790 clones in total (30 clones were analyzed per library); out of these, 1992 clones contained an SSU rDNA fragment and, subsequently, were sequenced. The BLAST search revealed that 1694 sequences had a high degree of similarity (97-100% identity) to sequences from AM fungal taxa and belonged to members of the phylum *Glomeromycota*. The rest of the sequences showed BLAST similarity to plants.

**Table 1.** Plant species growing as facultative epiphytes on date palm trees (Phoenix dactylifera) at the Historic Palm Grove of Elche (Alicante, Spain).

| Species | Abb | Life Form | Life Cycle | Origin |
| --- | --- | --- | --- | --- |
| *Asparagus acutifolius* L. | Aa | Woody | Perennial | Native |
| *Asparagus densiflorus* (Kunth) | Ad | Woody | Perennial | Exotic |
| *Asphodelus fistulosus* L. | Af | Herbaceous | Perennial | Native |
| *Brachychiton populneus* (Schott & Endl.) R. Br. | Bp | Woody | Perennial | Exotic |
| *Brachypodium distachyon* (L.) P. Beauv. | Bd | Herbaceous | Perennial | Native |
| *Centaurium spicatum* (L.) Fristch. | Cs | Herbaceous | Annual | Native |
| *Chenopodium murale* L. | Cm | Herbaceous | Annual | Native |
| *Conyza bonariensis* (L.) Cronquist | Cb | Herbaceous | Annual | Exotic |
| *Cynodon dactylon* (L.) Pers. | Cd | Herbaceous | Perennial | Native |
| *Euphorbia peplus* L. | Ep | Herbaceous | Annual | Native |
| *Jacaranda mimosifolia* D. Don | Jm | Woody | Perennial | Exotic |
| *Lantana camara* L. | Lc | Woody | Perennial | Exotic |
| *Medicago minima* (L.) L. | Mm | Herbaceous | Annual | Native |
| *Melia azedarach* L. | Ma | Woody | Perennial | Exotic |
| *Myrtus communis* L. | Mc | Woody | Perennial | Native |
| *Olea europea* L. | Oe | Woody | Perennial | Native |
| *Oxalis acetosella* L. | Oa | Herbaceous | Perennial | Exotic |
| *Oxalis pes-caprae* L. | Op | Herbaceous | Perennial | Exotic |
| *Parietaria judaica* L. | Pj | Herbaceous | Annual | Native |
| *Piptatherum miliaceum* (L.) Coss. | Pm | Herbaceous | Perennial | Native |
| *Pittosporum tobira* (Thunb.) W.T. Aiton | Pt | Woody | Perennial | Exotic |
| *Rhamnus alaternus* L. | Ra | Woody | Perennial | Native |
| *Rubia peregrina* L. | Rp | Herbaceous | Perennial | Native |
| *Sedum sediforme* (Jacq.) Grulich. | Sse | Herbaceous | Perennial | Native |
| *Setaria viridis* (L.) P. Beauv. | Sv | Herbaceous | Annual | Native |
| *Solanum nigrum* L. | Sn | Herbaceous | Annual | Exotic |
| *Sonchus oleraceus* L. | So | Herbaceous | Annual | Native |
| *Sonchus tenerrimus* L. | St | Herbaceous | Annual | Native |
| *Stenotaphrum secundatum* (Walter) Kuntze | Ss | Herbaceous | Perennial | Exotic |
| *Urtica urens* L. | Uu | Herbaceous | Annual | Native |
| *Veronica persica* Poir. | Vp | Herbaceous | Annual | Native |

Representative sequences of OTUs from root samples of epiphytic species were submitted to the GenBank database and are shown in bold in Fig. S1.

### Phylogenetic analysis of AMF groups

After phylogenetic analyses of the sequences, 23 AM fungal OTUs were detected in this study (Fig. S1; Table S1). Sequences of the families *Glomeraceae* (6 OTUs), *Diversisporaceae* (3 OTUs), *Gigasporaceae* (1 OTU), *Claroideoglomeraceae* (5 OTUs), *Paraglomeraceae* (7 OTUs), and *Archaeosporaceae* (1 OTU) were obtained. Eleven groups of AMF sequences or OTUs - namely *Glomus macrocarpum* (Glo1), *Septoglomus constrictum* (Sep), *Funneliformis mosseae-fragilistratum-caledonium-geosporum-coronatum* group (Fu), *Sclerocystis sinuosa* (Sc), *Rhizophagus intraradices-irregularis-fasciculatus* group (Rh), *Diversispora spurca-aurantia-eburnea* group (Div1), *Redeckera fulvum* (Red), *Scutellospora aurigloba-callospora* group (Scut), *Claroideoglomus luteum-claroideum-lamellosum-etunicatum* group (Cl2), *Paraglomus laccatum-occultum-brasilianum* group (Pa3), and *Archaeospora schenckii-trappei* group (Arch) - clustered with previously identified AMF sequences. Seven OTUs (Glo2, Div2, Cl3, Cl4, Cl5, Pa5, Pa7) did not cluster with any known *Glomeromycota* sequence. The remaining five OTUs (Cl1, Pa1, Pa2, Pa4, Pa6) were not related to any sequences of AMF in the database.

### Effect of plant origin, life-cycle, life-form and plant species on AM fungal community composition

In order to determine whether the number of clones sequenced was sufficient to represent the AM fungal diversity in all the facultative epiphytic species, rarefaction curves were constructed (Fig. 1). With the clones sequenced for each root sample, we covered perfectly the diversity of the AM fungal communities in all the plant species studied, since there was a well-defined leveling-off of all curves, and it is highly unlikely that the sequencing of more clones would have revealed more OTUs.

**Figure 1.**
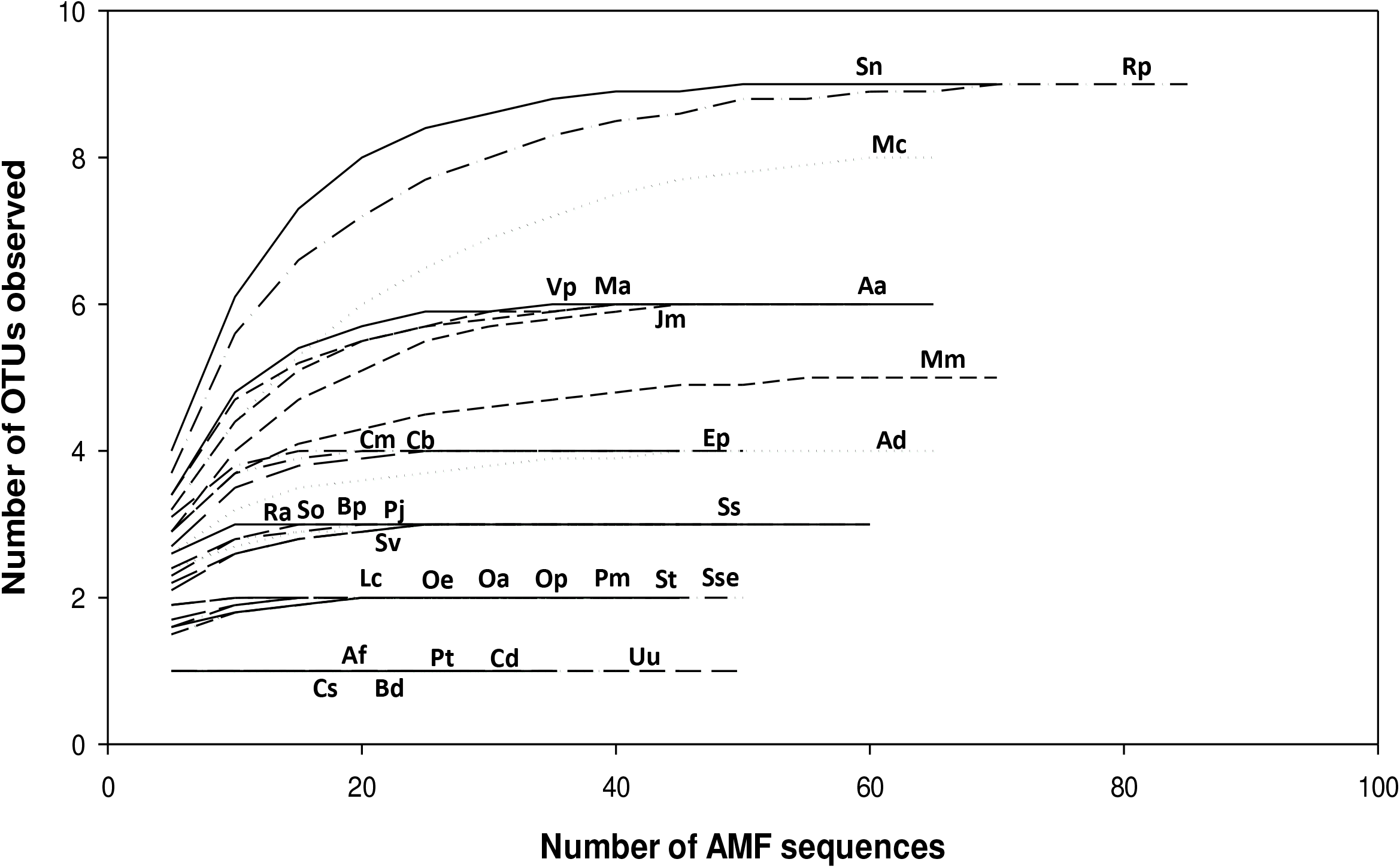
Sampling effort curves for the AM fungal community in the plants species growing as facultative epiphytes on date palm trees (*Phoenix dactylifera*). Aa: *Asparagus acutifolius;* Ad: *Asparagus densiflorus;* Af: *Asphodelus fistulosus;* Bp: *Brachychiton populneus;* Bd: *Brachypodium distachyon;* Cs: *Centaurium spicatum;* Cm: *Chenopodium murale;* Cb: *Conyza bonariensis;* Cd: *Cynodon dactylon;* Ep: *Euphorbia peplus;* Jm: *Jacaranda mimosifolia;* Lc: *Lantana camara;* Ma: *Melia azedarach;* Mc: *Myrtus communis;* Mm: *Medicago minima;* Oa: *Oxalis acetosella;* Oe: *Olea europea;* Op: *Oxalis pes-caprae;* Pj: *Parietaria judaica;* Pm: *Piptatherum miliaceum;* Pt: *Pittosporum tobira;* Ra: *Rhamnus alaternus;* Rp: *Rubia peregrina;* Sn: *Solanum nigrum;* So: *Sonchus oleraceus;* Sse: *Sedum sediforme;* Ss: *Stenotaphrum secundatum;* St: *Sonchus tenerrimus;* Sv: *Setaria viridis;* Uu: *Urtica urens;* Vp: *Veronica persica*.

An indicator species analysis (ISA) was conducted to find specific OTUs associated with plant origin, life-cycle, and life-form (Table 2). When the ISA was performed considering the plant origin as the grouping factor, two OTUs were associated with the exotic group (Cl4 and Pa7) and one OTU with the native group (Glo2). Three OTUs were found to be specific for the roots of annual plants when the plant life-cycle factor was considered (Glo1, Fu, Glo2). With respect to plant life-form, one OTU was specific for woody species (Pa7).

**Table 2.** Indicator species analyses.

| OTUs associated to plant origin |  |  |  |  |
| --- | --- | --- | --- | --- |
| Group Exotic |  |  |  |  |
| #sps. 1 | A | B | Index | p-value |
| Cl4 | 0.8791 | 0.2424 | 0.462 | 0.0035 |
| Pa7 | 1.0000 | 0.0909 | 0.302 | 0.0473 |
| Group Native |  |  |  |  |
| #sps. 1 |  |  |  |  |
| Glo2 | 1.0000 | 0.1667 | 0.408 | 0.0307 |
| OTUs associated to plant life cycle |  |  |  |  |
| Group Annual |  |  |  |  |
| #sps. 4 |  |  |  |  |
| Glo1 | 0.5746 | 0.8056 | 0.680 | 0.0297 |
| Fu | 0.7037 | 0.3333 | 0.484 | 0.0075 |
| Glo2 | 0.8636 | 0.2222 | 0.438 | 0.0063 |
| OTUS associated to plant life form |  |  |  |  |
| Group Woody |  |  |  |  |
| #sps. 1 |  |  |  |  |
| Pa7 | 1.0000 | 0.1000 | 0.316 | 0.0323 |

As shown by the perMANOVA, the plant species factor had highly significant effects on the AMF communities composition and structure (F=14.45, P=1e-04). Also, the lifecycle and life-form significantly influenced the distribution of the AMF (F=2.4082, P=0.0365 and F= 3.0367, P=0.0128, respectively), whereas the plant origin did not have a significant effect (F=1.6434, P=0.1250) (Table 3).

**Table 3.** PERMANOVA analysis of the effect of plant origin, life cycle, life form and plant species on the distribution of AMF OTUs in plant species growing as facultative epiphytes on date palm trees (Phoenix dactylifera) under semiarid Mediterranean conditions.

|  | Df | SS | MS | F. Model | R<sup>2</sup> | Pr (>F) |
| --- | --- | --- | --- | --- | --- | --- |
| Origin | 1 | 0.406 | 0.406 | 1.643 | 0.017 | 0.125 |
| Life cycle | 1 | 0.594 | 0.595 | 2.408 | 0.025 | **0.036\*** |
| Life form | 1 | 0.750 | 0.750 | 3.037 | 0.032 | **0.013\*** |
| Plant species | 30 | 20.75 | 0.692 | 14.46 | 0.874 | **1e-04\*\*\*** |

Df, degrees of freedom; SS, sum of squares; MS, mean squares; Pr value by permutation
\*In bold, statistically significant relationships ( $P \leq 0.05$ )

## DISCUSSION

In this study we found a high number of facultative epiphytic plant species growing on *Phoenix dactylifera* palm trees being colonized by arbuscular mycorrhizal fungi under semiarid Mediterranean conditions (Table 1). There is only a recent study carried out with one plant species, *Sonchus tenerrimus* L. growing as facultative epiphyte in *P. dactylifera* in semiarid conditions (44), the rest of studies concerning the occurrence of AM symbiosis in epiphytic vascular plants have been conducted in temperate and tropical ecosystems with plant species such as ferns and lycophytes (15,27,32,50).

In our study the AMF richness was covered in the thirty one epiphytic plants studied with the method used, since all accumulation curves reached a well-defined asymptote. Consistent results for the fungal community structure and relative abundances of fungal taxa can be obtained whatever the sequencing approach used; i.e., NGS vs cloning-Sanger sequencing (29).

The species indicator analysis revealed three OTUs (Glo1, Glo2 and Fu) belonging to the *Glomeraceae* and associated with annual species, Glo2 was also an indicator taxon for native species. One OTU (Pa7) belonging to the *Paraglomeraceae* was an indicator for both woody and exotic species (Table 3). These OTUs showed similarity to database sequences reported previously from a wide range of environmental conditions in terrestrial habitat, varying from stressful to optimal conditions (1,14,18,42,48,49). This shows the high plasticity of these AMF taxa.

Epiphytic habitats are characterized by nutritional insufficiency (3), since the substrate on which epiphytic plants grow is accumulate organic matter and rainwater. In our case, the different facultative epiphytic plants species grew in small ledges of cut leaves in the trunks of date palms (*P. dactylifera*) with high organic carbon content (36%) and subjected to large periods of low water availability. Under these conditions, it has been suggested a set of morphological and physiological adaptations in epiphytic taxa to cope water and nutrient shortage (25,36), being reported only the association with arbuscular mycorrhizal fungi in Mediterranean semiarid ecosystems (44).

Our results suggest that the AMF community composition is plant host identity-dependent. Plant affiliation to a species strongly influenced the AMF assemblages in the epiphytic environment (Table 3). This agrees with the hypothesis put forward to explain the phylogenetic distribution of mycorrhizas in land plants (46) and suggests that plant phylogeny might influence the AMF community - as proposed by Montesinos-Navarro, et al. (24), who demonstrated that changes in the phylogenetic composition of plant and AMF assemblages do not occur independently in a patchy environment. The differences in dispersal processes between AMF and plants might explain which group drives the other. According to this hypothesis, in primary successions plants arrive before AMF and then act as a potential filter for the latter, as seems to be the case in the AMF community in palm trunks.

Functional traits such as plant life-form and plant cycle also shaped the AMF interactions. The AMF community differed between the annual and perennial epiphytic species (F=2.4082; *P*=0.0365) and between herbaceous and woody plant species (F=3.0367; *P*=0.0128). The dependence of the composition of the AMF community on the host-plant has been previously reported in different habitats around the world (1,2,6,17,39,42), with different plant ecological groups (e.g. habitat generalists vs specialists) (7,43) or ecosystems (45). In a study in the same semiarid Mediterranean conditions, Alguacil et al. (2) found a different AMF community composition and greater diversity in perennial plant species than in annuals, in accordance with our results (where we detected greater AMF richness in perennial (21 OTUs) than in annual plants (15 OTUs)). These authors explained these differences as being due to the fact that the former are in the soil for longer, giving more opportunities for mycorrhization establishment. Similarly, the continuity of the roots of an individual with time is critical in an epiphytic environment, taking into account the strong limitations to AMF dispersal.

Several authors have pointed out the constraints to AMF dispersal in epiphytic habitats, as propagules are mainly dispersed by biotic vectors (10,21,44). Moreover, in this particular habitat, plants generally grow as individuals inside the cut leaves without any connections among root systems, thus preventing AMF spread. Under these circumstances spatial and temporal coincidence of the plants and fungi, which is higher in perennials plants than in annuals, is crucial for mycorrhiza development.

In this work the epiphytic plants associated with AMF irrespective of whether they are native or not. This is in accordance with Bunn, et al. (4), who compared 67 publications with divergent hypotheses about the enhancement or loss of AMF following invasions by exotic plants and concluded that other factors - such as plant functional groups - better predict AM variations in fungal associations (not only in terms of AM status but also fungal colonization or growth response), as we also found for non-invasive exotic species. Unlike invasive plants - which have specific functional traits that confer the ability to colonize new habitats, usually displacing native species - no invasive ability was observed in the exotic plant species growing in the epiphytic environment provided by the palm orchards.

In conclusion, a high diversity of facultative epiphytic plant species are colonized by AMF in the semiarid Mediterranean conditions studied. Plant identity and the functional plant life-cycle and life-form features determine the AMF assemblages.

## MATERIAL AND METHODS

### Study area and sampling

The study area was located at the Historic Palm Grove of Elche (Alicante), southern Spain (38°15’ 51, 28’’N, 0° 41’ 51, 23’’ W, 82 m.a.s.l). It covers 550 Ha and contains about 180.000 adult, *Phoenix dactylifera* date palms aged 40-60, planted in one or two rows around cultivated, rectangular-shaped orchard (35). The climate is Mediterranean semiarid, with a mean annual temperature of 17.7°C and a mean annual rainfall of 266 mm (www.aemet.es). Three sampling plots separated at least 300 m from each other, and each consisting of two palm orchards (2.000 m^2^ area, 250 palm trees Ha^-1^), were selected. A total of 31 plant species growing epiphytically between 1.5-2 m height on date palm trunks were sampled at the spring growing season (three individuals of each species, one per plot). Plant species were classified according to their origin into native or exotic (authoctonous or aloctonous), according to their life cycle into annual or perennial and according to life form into herbaceous or wood. Canopy or bare-limb epiphytes were not found. Roots systems were collected, fine foots were separated, briefly rinsed, quickly dried on paper and used for molecular analysis.

### Roots DNA extraction and PCR

For each sample, 0.2 g fresh root material was frozen with liquid nitrogen, placed into a 2-ml screw-cap propylene tube together with two tungsten carbide balls (3 mm) and ground (3 min, 13000 r.p.m.) using a mixer mill (MM 400, Retsch, Haan, Germany). Total DNA was extracted using a DNeasy Plant Mini Kit following the manufacturer’s recommendations (Qiagen). The extracted DNA was resuspended in 20 μl of water and stored at −20°C.

Several dilutions of extracted DNA (1/10, 1/50, 1/100) were prepared and 2 μl were used as template. Partial small subunit (SSU) ribosomal RNA gene fragments were amplified using nested PCR with the universal eukaryotic primers NS1 and NS4 (47). PCR was carried out in a final volume of 25 μl using PuReTaq^™^ Ready-To-Go PCR beads (Amershan Pharmacia Biotech), 0.2μM dNTPs and 0.5 μM of each primer (PCR conditions: 94 °C for 3 min, then 30 cycles at 94 °C for 30 s, 40 °C for 1 min, 72 °C for 1 min, followed by a final extension period at 72 °C for 10 min).

Then, 2μ1 from the first PCR were used as template DNA in a second PCR reaction performed using the specific primers AML1 and AML2 (16). PCR reactions were carried out in a final volume of 25 μl using the PuReTaq™ Ready-To-Go PCR beads (Amershan Pharmacia Biotech), 0.2 μM dNTPs and 0.5 μlM of each primer (PCR conditions: 94 °C for 3 min, then 30 cycles of 1 min denaturation at 94 °C, 1 min primer annealing at 50 °C and 1 min extension at 72 °C, followed by a final extension period of 10 min at 72 °C). Positive and negative controls using PCR positive products and sterile water respectively were also included in all amplifications. All the PCR reactions were run on a Perkin Elmer Cetus DNA Thermal Cycler. Reactions yields were estimated by using a 1.2% agarose gel containing *GelRed*^™^ (Biotium).

### Cloning and sequencing

The PCR products of the expected band length, approximately 795 bp were purified using a Gel extraction Kit (Qiagen) cloned into pGEM-T Easy vector (Promega) and transformed into *Escherichia coli* (Xl1 blue). Putative positive transformants were screened in each resulting SSU rRNA gene library, using 0.7 unit of RedTaq DNA polymerase (Sigma) and the supplied reaction buffer to a final volume of 25μ and a reamplification with AML1 and AML2 primers with the same cycling conditions described above. Product quality and size were checked in agarose gels as described above. All clones having inserts of the correct size (795 bp) in each library were sequenced using the universal primers SP6 and T7 by Laboratory of Sistemas Genómicos (Valencia, Spain).

### Phylogenetical analysis

Sequence editing was done using the program FinchTV 1.4.0 (Geospiza, Inc.; Seattle, WA, USA; http://www.geospiza.com). A search for similar sequences to the ones from this study was conducted with the BLAST tool (51) provided by GenBank. Phylogenetic analysis was carried out on the sequences obtained in this study and those corresponding to the closest matches from GenBank as well as sequences from cultured AMF taxa including representatives of the major taxonomical groups described by (34). All the sequences were aligned, using the multiple sequence comparison program, MAFFT, version 7.0 (available at http://align.bmr.kyushu-u.ac.jp/mafft/software) and the alignment was adjusted manually in BioEdit software version 7.2.5. (12). The program CHIMERA_CHECK 2.7 (Ribosomal Database Project II; http://rdp.cme.msu.edu) was used to check for chimeric artifacts among the 18S rDNA sequences.

Maximum likelihood (ML) phylogenetic tree inference was performed with MEGA software (version 5.05) (40). Nucleotide data files were first tested to find the best DNA evolution model. The general time reversible model with a discrete gamma distribution showed the lowest Bayesian information criterion (BIC) scores and was deemed to best describe the nucleotide substitution pattern. Initial trees for the heuristic search were obtained by applying the neighbor-joining method to a matrix of pairwise distances estimated using the maximum composite likelihood (MCL) approach. The robustness of all trees obtained was evaluated by 1000 bootstrap replications. *Endogone pisiformis* Link and *Mortierella polycephala* Coem, were used as the out-groups.

Different AMF sequence types or OTUs (operational taxonomic units), were defined as groups of closely related sequences, with a high level of bootstrap support in the phylogenetic analyses (higher than 80%) and sequence similarity ≥ 97%.

### Statistical analysis

The number of clones for each AM fungal OTUs in each plant species was used to calculate the rarefaction curves. The rarefaction curves were produced by plotting the number of OTUs observed against the number of sequences obtained using the freely available Analytic Rarefaction software (version 1.3) (http://www.uga.edu/~strata/software/anRareReadme.html).

In order to ascertain whether the AMF communities composition and structure were significantly affected by the experimental factors (plant origin, life form, life cycle and plant species) a permutational multivariate analysis of variance (perMANOVA) was performed with the adonis function in vegan (28) using the Jaccard distance matrix and 999 permutations.

Three different indicator species analyses (ISA) were conducted on the OTUs presence/absence dataset using the “indicspecies” package implemented in R (8). This analysis allows us identifying species which are associated to a classifier factor by calculating an Indicator Value (IndVal) (9). We considered as classifier factors the plant origin, life form and life cycle. The statistical significance of the indicator values was tested using a permutation test with 999 permutations.

The AMF OTUs richness was subjected to ANOVA to test for significant differences between different factors (plant origin, life cycle, life form and plant species). For the comparisons among means the Duncan’s test at P<0.05 was used. All the statistical procedures were carried out with the software package IBM SPSS Statistic 24.0 for Windows.

#### Nucleotide sequence accession numbers

A total of 109 representative sequences of OTUs from root samples generated in this study have been deposited at the National Centre for Biotechnology Information (NCBI) GenBank (http://www.ncbi.nlm.nih.gov) under the accession numbers MG835459-MG835567.

## AKNOWLEDGEMENTS

MM Alguacil was supported by the Ramon and Cajal programme (Ministerio de Educación y Ciencia, Spain). This research was supported by the Spanish Plan Nacional-FEDER Projects CGL-2015-64168-R and CGL2013-42312-R.

The authors declare that the research was conducted in the absence of any commercial or financial relationships that could be construed as a potential conflict of interest.

